# Glucose-dependent regulation of hepatic adipsin controls glucose uptake and tolerance

**DOI:** 10.64898/2026.07.02.735968

**Authors:** Sujay Krishna Maity, Asmita Bhar, Abhishek Sen, Tanusree Das, Atin Sasmal, Souveek Mitra, Abhijit Chowdhury, Partha Chakrabarti

## Abstract

Complement factor D, also known as adipsin, is produced by adipose tissue, and the liver that links metabolic regulation with innate immunity. Despite its established systemic functions, the regulation of hepatic adipsin expression and its contribution to metabolic disease remain poorly defined. Here, we show that hepatic adipsin protein abundance is markedly increased in individuals with type 2 diabetes (T2D), and positively correlates with glycated hemoglobin, despite unchanged mRNA expression. Concordantly, hepatic adipsin protein levels were elevated in multiple murine models of hyperglycemia, including type 1 diabetes (T1D), T2D, and following fasting–refeeding transitions. In cultured hepatocytes, glucose exposure induced a rapid, dose-dependent increase in adipsin protein without altering transcript abundance, demonstrating post-transcriptional regulation. Mechanistically, glucose stimulates adipsin translation via dephosphorylation of eukaryotic initiation factor 2α (eIF2α), and activation of the mammalian target of rapamycin, mediated by the 5′ untranslated region of adipsin mRNA. Functionally, hepatocyte-specific depletion of adipsin impaired postprandial glucose tolerance, with reduced glucose uptake and a marked downregulation of glucose transporter type 2 (GLUT2). Taken together, these findings identify hepatic adipsin as a glucose-responsive translational target that couples nutrient availability to metabolic adaptation, revealing a new layer of regulation with potential relevance to diabetes pathogenesis.

**Highlights:**

- Hepatic adipsin protein increases in type 2 diabetes and correlates with glycemic status independent of mRNA expression.
- Glucose induces adipsin translation through eIF2α dephosphorylation and mTOR activation.
- mTOR controls adipsin synthesis via structured 5′UTR of adipsin mRNA.
- Liver-specific adipsin depletion impairs post-prandial glucose tolerance by downregulating GLUT2.
- Hepatic adipsin acts as a glucose-responsive effector of glycemic control.

## 1. Introduction

Adipsin, or CFD, resides at the nexus of metabolic regulation [1], immune function [2], and endocrine signaling pathways [3]. Adipsin is a serine protease [4] that catalyzes the rate-limiting step in the alternative complement pathway and has lately been shown to have biological significance beyond classical immunology [5]. Accumulating studies over the past decade have identified it as a metabolically active protein [6] that is predominantly secreted by adipose tissue [7; 8] and macrophages [9]. Adipsin has been shown to be instrumental in the regulation of systemic inflammation [10], systemic energy balance [11], and chronological aging [12], suggesting that this dual involvement of adipsin, both in innate immunity and metabolic homeostasis, positions it as a crucial link between the complement pathway and metabolic disease pathogenesis.

The liver, an immunologically complex and central metabolic organ produces up to 90% of the fluid-phase complement proteins [13]. Adipsin, however, often believed not to be significantly produced nor secreted by hepatocytes. However, immunoblot analysis from primary culture of human hepatocytes [14], and a human hepatocyte cell line, HepG2 [15] exhibited that adipsin had a similar molecular weight compared to the factor D from normal human serum and U937 cells [16]. Additionally, recent studies have shown adipsin expression is altered during alcohol associated liver disease (ALD) [17], and metabolic dysfunction-associated liver disease (MASLD) [18]. These observations highlight multifaceted and is context dependent role of adipsin in liver. Several cross-sectional studies have reported elevated serum adipsin levels in patients with MASLD [19], correlating with metabolic burden and hepatic steatosis severity. However, paradoxically, other studies have shown inverse associations between adipsin levels and NAFLD risk, particularly in obese populations, and this association remained significant after adjusting for multiple metabolic confounders [20]. A prospective clinical study of a 3-year follow-up also showed that higher baseline adipsin levels were independently associated with lower odds of NAFLD remission [21]. Taken together, serum adipsin might serve as a biomarker in MASLD. The tissue-specific sources of adipsin, especially the liver, might have a distinct role that is not completely understood by measuring circulating levels alone.

In this study, we aim to explore the expression pattern of adipsin in both human and mice liver during various metabolic diseases and to elucidate the molecular mechanisms governing its regulation. Furthermore, we sought to define the functional role of liver-derived adipsin during hepatic metabolic perturbations. Our findings establish that hepatic adipsin is regulated by glucose through eIF2α- and mTOR-dependent translational control. Functionally, adipsin promotes glucose uptake via GLUT2, and its depletion impairs glycemic control. Together, these data define a glucose–eIF2α–mTOR–adipsin feedback loop linking nutrient sensing to hepatic glucose metabolism.

## 2. Results

### 2.1 Adipsin protein expression is increased in the livers of T2D patients and diabetes mouse models

Human hepatocytes express adipsin [15; 16]; however, the factors regulating hepatic adipsin expression are unknown. We assessed adipsin protein and mRNA expressions in consecutive liver biopsy samples from MASLD patients with (n=10) or without (n=20) diabetes. NAS, fibrosis scores, and BMI were comparable between the groups, whereas glycemic parameters such as FBS and HbA1c were expectedly higher in patients with diabetes (Table 1). Although we did not find a difference in adipsin gene expressions (Figure 1A), protein levels determined by ELISA of tissue homogenate were significantly higher in diabetes patient livers (Figure 1B). Adipsin protein identified by immunohistochemistry also revealed enhanced expression in diabetes patients (Figure 1C). Interestingly, pericentral stain was more prominent and Kupffer cells had the maximum intensity. Next, we performed correlation analysis to identify how liver adipsin is associated with the parameters of glycemia and MASLD. Only HbA1c was found to be significantly correlated with adipsin protein levels (r=0.53, p=0.003), suggesting long-term hyperglycemia could induce hepatic adipsin (Figure 1D). In contrast to liver adipsin, we did not find a significant difference in plasma adipsin levels between the groups (Figure 1E). We next examined whether hyperglycemia and its therapy could modulate liver adipsin levels in experimental animal models of T1D and T2D. First, we used streptozotocin (STZ) to induce T1D, followed by insulin treatment (0.75U/kg) for seven days [22] (Figure 1F). Whilst STZ caused rapid hyperglycemia, insulin therapy normalized it (Figure 1G). Body and liver weight, and serum adipsin levels were unaltered by insulin treatment (Figure 1H-J). Adipsin gene expression in liver showed no significant change following STZ or insulin administration (Figure 1K). In contrast, adipsin protein levels were intriguingly enhanced following STZ injection, and was markedly diminished by insulin treatment (Figure 1L). IHC also revealed a consistent pattern of adipsin expression (Figure 1M). In a separate experimental design, a T2D model was established through high fat diet (HFD) feeding for 12 weeks (Supplementary Figure 1A). HFD-fed mice exhibited increased body weight, hepatic mass, and epididymal adipose tissue mass, accompanied by elevated serum triglycerides, cholesterol, and hepatic transaminases (AST and ALT) (Supplementary Figure 1B–H). Fasting hyperglycemia and elevated plasma insulin concentrations in HFD-fed mice confirmed the development of insulin resistance (Figure 2N, O). Notably, these mice also displayed increased circulating adipsin levels (Figure 1P). Hepatic adipsin mRNA expression in these animals was variable and did not show a consistent trend (Figure 1Q); however, hepatic adipsin protein content was significantly increased (Figure 1R, S). Collectively, these data indicate that hyperglycemia per se induces upregulation of hepatic adipsin protein expression in murine models of diabetes.

**Figure 1.**
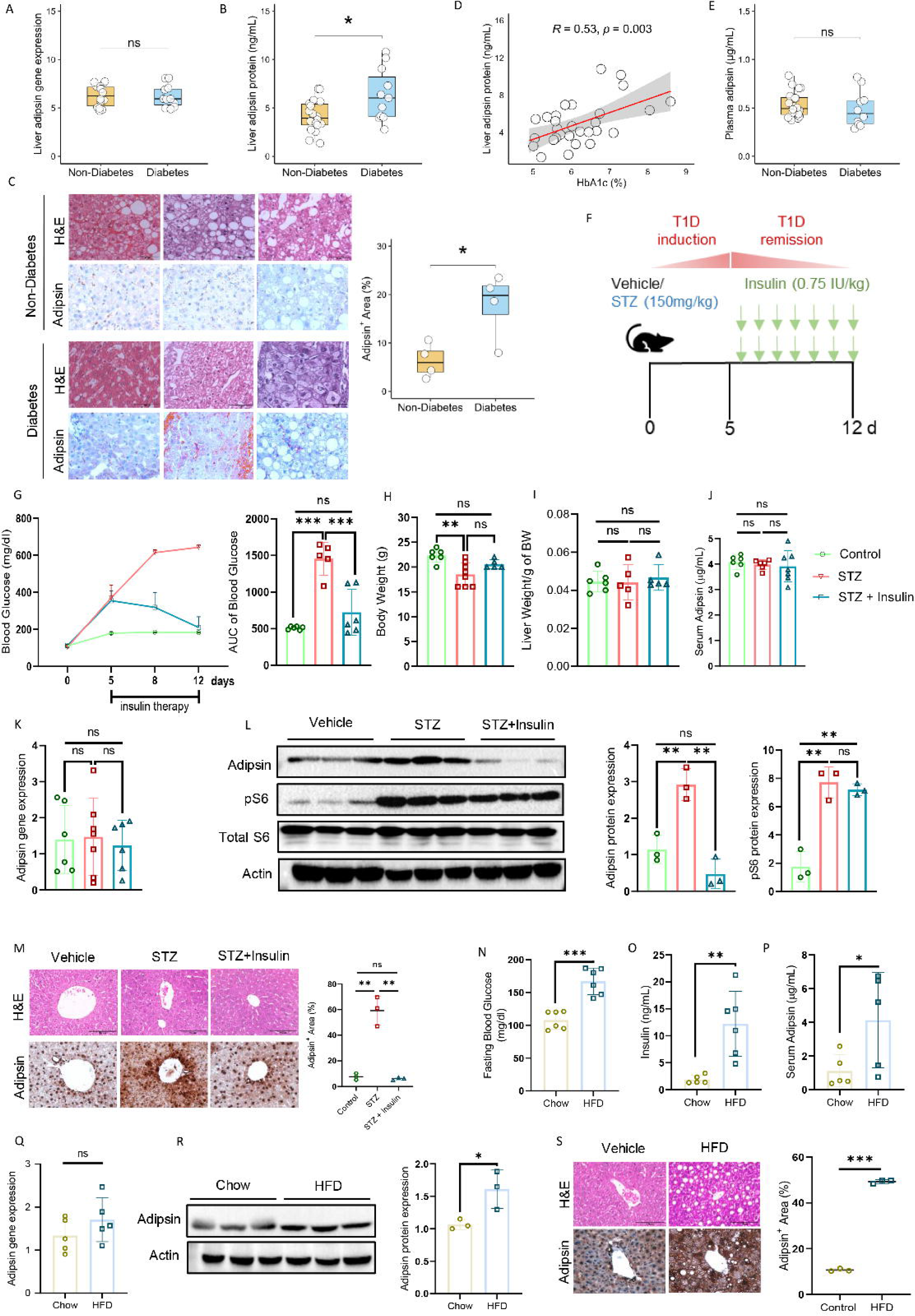
Hepatic adipsin protein expression is elevated in T2D patients, murine models of diabetes and correlates with chronic hyperglycemia. (A) Hepatic adipsin mRNA expression in non-diabetes (n = 10) and diabetes (n = 20) MASLD patients. (B) Adipsin protein levels in liver homogenates from MASLD patients with or without diabetes. (C) Left panels: H&E, immunohistochemistry of adipsin in independent liver biopsy. Right panels: Hepatic adipsin positive area (%) was quantified using ImageJ software (n=4/group, 6 fields/sample). (D) Correlation between hepatic adipsin protein and HbA1C. (E) Plasma adipsin concentrations in non-diabetes and diabetes patients. (F) Experimental design for STZ-induced type 1 diabetes (T1D) followed by glycemic correction using insulin treatment. (G) Blood glucose levels and corresponding area under the curve (AUC). (H–J) Body weight, liver weight, and serum adipsin concentrations. (K) Hepatic adipsin mRNA expression in control, STZ-treated, and STZ plus insulin-treated mice. (L) Immunoblot of adipsin, phosphorylated S6 (pS6), and total S6 in liver homogenates. Right panel: Densitometric analysis of adipsin and pS6 (n=3). (M) H&E, immunohistochemistry of adipsin on liver sections. Hepatic adipsin positive area (%) quantified (n=3/group, 6 fields/sample). (N–O) Fasting blood glucose and plasma insulin concentrations in the high fat diet (HFD)-induced type 2 diabetes (T2D) model. (P) Circulating adipsin concentrations in HFD-fed mice. (Q) Hepatic adipsin mRNA expression. (R) Immunoblot of adipsin from liver homogenates. Bottom panel: Densitometric analysis of adipsin (n=3) (S) H&E, immunohistochemistry of adipsin on liver sections. Mice n = 6 (control), n = 8 (STZ), n = 7 (STZ + insulin), n= 6 (Chow), n=6 (HFD). Data are presented as mean ± SD. *\*P*< 0.05, *\*\*P*< 0.01, *\*\*\*P*< 0.001, statistical significance was determined using the Mann–Whitney U test, Spearman correlation analysis was performed to assess associations between variables, and unpaired two-tailed t-test or one-way ANOVA followed by Bonferroni’s multiple comparison test, as appropriate. ns, not significant; GGT, γ-glutamyl transferase; STZ,streptozotocin; HFD, High Fat Diet; BW, body weight. Scale bar, 100 μm.

**Figure 2.**
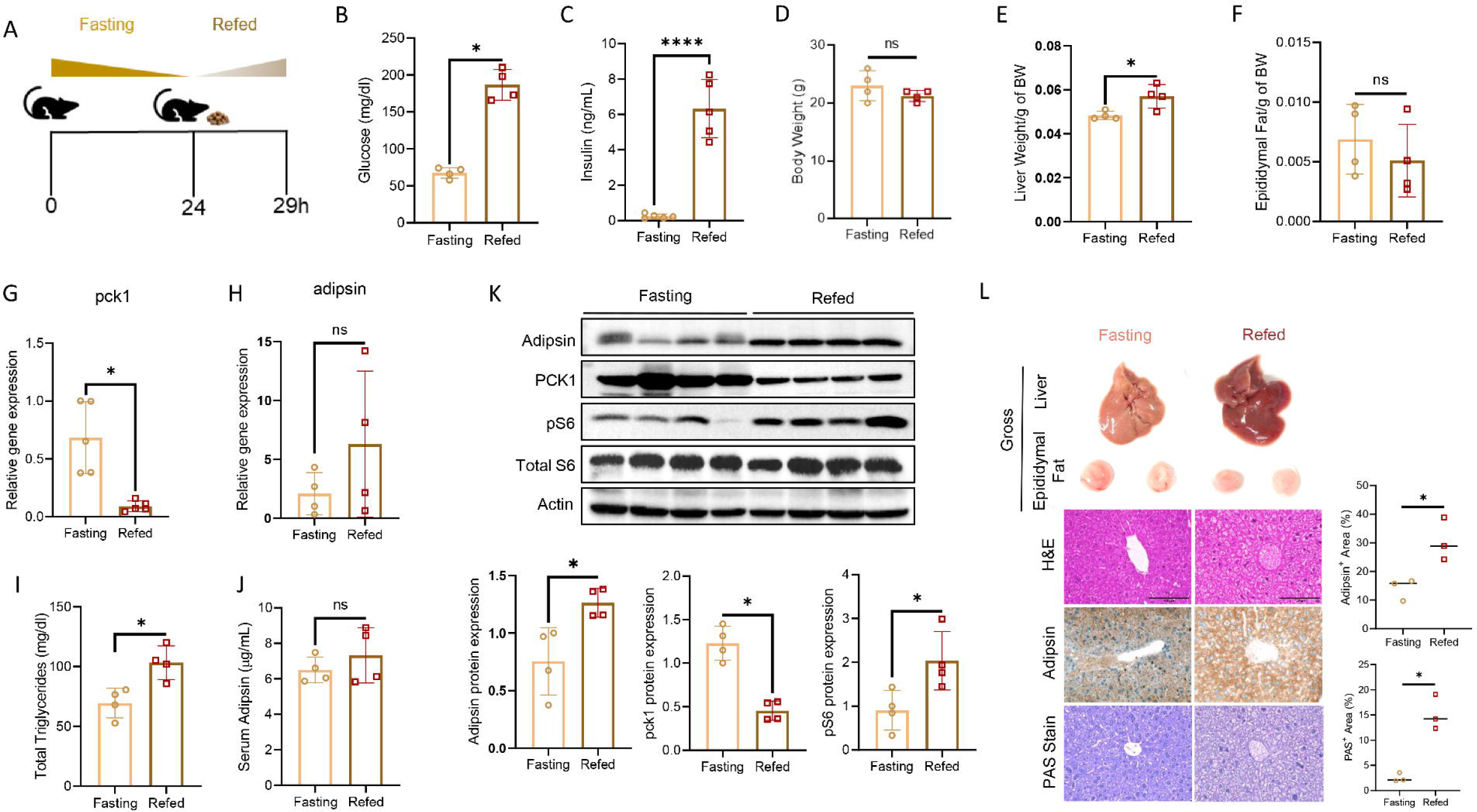
Regulation of hepatic adipsin expression in fasting-refeeding cycle. (A) Experimental design of 24-hour fasting followed by 5-hour refeeding. (B–C) Plasma glucose and insulin concentrations during fasting and refeeding. (D–F) Body weight, liver weight, and epididymal adipose tissue weight during fasting and refeeding. (G–H) Hepatic adipsinand PCK1 mRNA expression during fasting and refeeding. (I–J) Plasma triglyceride and circulating adipsin concentrations during fasting and refeeding. (K) Immunoblots of hepatic adipsin and PCK1 in liver homogenates. Bottom panels: Densitometric analysis of adipsin, PCK1 and pS6 levels (n=4). (L) Gross appearance of liver and epididymal adipose tissue depot. H&E, immunohistochemistry and Periodic Acid-Schiff (PAS) on liver sections. For immunohistochemistry, hepatic adipsin-positive area (%) was quantified from representative sections (n = 3 mice per group, 6 fields per sample). Mice n = 8 (fasting), n = 8 (refed)). Data are presented as mean ± SD. *\*P*< 0.05, *\*\*P*< 0.01, *\*\*\*P*< 0.001, statistical significance was determined using unpaired two-tailed t-test. ns, not significant; BW, body weight Scale bar, 100 μm.

**Table 1.** Baseline clinical and biochemical characteristics of study participants.

|  | Non-Diabetes<br>(n=20) | Diabetes<br>(n=10) | P-value |
| --- | --- | --- | --- |
| Sex (Male/Female) | 11/9 | 7/3 | 0.4497 |
| Age (Years) | 38.93 ± 10.86 | 41.55 ± 6.55 | 0.3320 |
| BMI (kg/m<sup>2</sup>) | 28.54 ± 5.01 | 31.65 ± 5.26 | 0.1200 |
| Fasting Glucose (mmol/L) | 5.38 ± 0.73 | 6.32 ± 1.41 | 0.0402 |
| HbA1c (%) | 5.52 ± 0.41 | 7.15 ± 0.66 | < 0.0001 |
| Fasting Insulin (μIU/mL) | 11.76 ± 4.33 | 15.42 ± 7.95 | 0.2230 |
| ALT (U/L) | 70.07 ± 33.11 | 68.5 ± 35.85 | 0.9500 |
| AST (U/L) | 56.99 ± 25.39 | 52.55 ± 22.85 | 0.6170 |
| GGT (U/L) | 34.58 ± 27.73 | 46 ± 31.84 | 0.1173 |
| Total Triglycerides (mg/dL) | 176.04 ± 81.99 | 201.4 ± 63.16 | 0.1280 |
| Total Cholesterol (mg/dL) | 187.65 ± 45.79 | 187.65 ± 45.79 | 0.6640 |
| NAFLD Activity Score | 2.58 ± 1.47 | 1.9 ± 1.2 | 0.1350 |
| Steatosis score | 1.58 ± 0.9 | 1.2 ± 0.79 | 0.1610 |
| Fibrosis score | 1.23 ± 1.21 | 1.7 ± 1.42 | 0.3300 |
| HOMA-IR | 3.43 ± 1.39 | 5.53 ± 3.74 | 0.1970 |
| HOMA-B | 138.56 ± 59.92 | 115.38 ± 58.35 | 0.2970 |

### 2.2 Fasting refeeding cycle modulates adipsin protein levels in mice liver

We investigated whether postprandial elevations in blood glucose exert a similar regulatory effect on hepatic adipsin expression. For this purpose, mice were subjected to a 24 h fasting period [23], followed by refeeding with a standard chow diet for 5 h (Figure 2A). As anticipated, refeeding resulted in a pronounced increase in circulating glucose and insulin concentrations (Figure 2B, C), accompanied by a modest elevation in liver mass, without appreciable changes in body weight or epididymal adipose tissue weight (Figure 2D-F). Hepatic expression of the rate-limiting gluconeogenic gene PCK1 was markedly suppressed under the re-fed condition, whereas adipsin mRNA abundance remained unaltered (Figure 2G, H). Plasma triglycerides level increased while plasma adipsin concentrations were unaffected by the fasting–refeeding transition (Figure 2I, J). Immunoblotting further revealed a striking drop in hepatic PCK1 protein in the re-fed state, in contrast to a significant elevation in hepatic adipsin protein levels (Figure 2K). Consistently, histological and immunohistochemical analyses showed a significant increase in both adipsin levels and glycogen content in the fed state (Figure 2L). Collectively, these findings indicate that hepatic adipsin protein abundance is increased not only under pathological hyperglycemia associated with diabetes but also during physiological fluctuations in glucose availability.

### 2.3 Glucose controls adipsin protein translation via eIF2α and mTOR

Because hyperglycemia increased hepatic adipsin expression, whereas adipsin depletion impaired glucose tolerance, we next asked how glucose availability regulates adipsin production. Given that glucose deprivation induced eIF2α phosphorylation and suppressed mTOR signalling [24; 25] (Figure 3A), suggesting that adipsin expression might be controlled at the level of translation through coordinated regulation of these nutrient-sensing pathways.To formally test whether adipsin is subject to cell-autonomous translational control, primary hepatocytes and HepG2 cells were transiently deprived of media and subsequently re-exposed to increasing concentrations of glucose. Media withdrawal led to a rapid decline in adipsin protein levels (Figure 3B; Supplementary Figure 2A), which were restored in a dose-dependent manner upon glucose reintroduction (Figure 3C; Supplementary Figure 2B, D), without changes in mRNA abundance (Supplementary Figure 2C), indicating post-transcriptional regulation. Consistently, glucose deprivation enhanced eIF2α phosphorylation and reduced mTORC1 activity, as evidenced by diminished 4E-BP1 phosphorylation (Figure 3B). Glucose supplementation reversed these effects, suppressing eIF2α phosphorylation while augmenting 4E-BP1 phosphorylation (Figure 3C). To determine whether glucose metabolism is required for adipsin induction, cells were pre-treated with 2-deoxy-D-glucose (2-DG), a glycolytic inhibitor. 2-DG markedly attenuated glucose-induced adipsin protein expression, concomitant with increased eIF2α phosphorylation and reduced 4E-BP1 phosphorylation (Figure 3D), demonstrating that metabolic flux, rather than glucose sensing alone is necessary. Given that insulin activates the AKT–mTOR axis, we next assessed whether insulin alone is sufficient to induce adipsin. Interestingly, insulin treatment in primary hepatocytes did not increase adipsin expression (Supplementary Figure 2D).Although insulin robustly activated AKT and mTOR signaling in the absence of glucose, it failed to restore adipsin protein expression (Figure 3E). Notably, insulin did not suppress eIF2α phosphorylation under glucose deprivation. In vivo, insulin administration under fasting conditions reduced blood glucose (Figure 3F), and activated hepatic AKT–mTOR signaling without altering eIF2α phosphorylation or adipsin expression (Figure 3G). Similarly, in STZ-diabetic mice, restoration of blood glucose by insulin treatment (Figure 1G) caused an increase in eIF2α phosphorylation (Supplementary Figure 2G).Collectively, these data demonstrate that glucose availability is both necessary and sufficient to induce hepatocyte adipsin expression through coordinated regulation of the eIF2α and eIF4E translational nodes. In contrast, insulin selectively activates the mTOR arm but is insufficient to drive adipsin synthesis in the absence of glucose, underscoring a dominant role for metabolic control of translation in governing hepatic adipsin expression.

**Figure 3.**
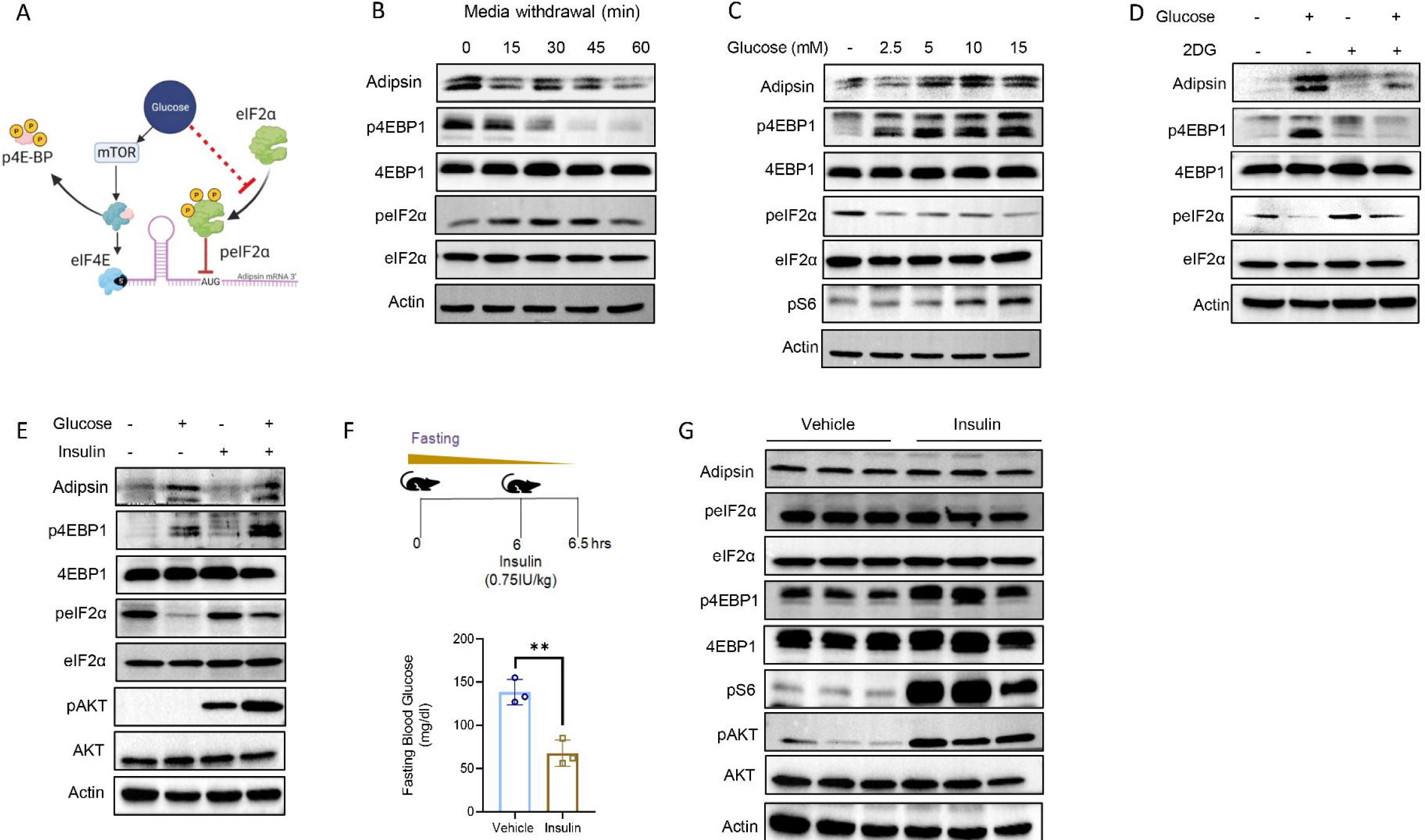
Glucose regulates hepatic adipsin protein translation via eIF2α and mTOR. (A) Experimental hypothesis illustrating that glucose availability is required for hepatic adipsin protein expression through coordinated regulation of peIF2α and eIF4E-dependent translation. (B-C) Immunoblots from HepG2 cells following media withdrawal for indicated time points (B), and after glucose re-addition in a dose dependent manner (C). HepG2 cells were starved in HBSS for 6h prior to treatments. (D) Cells were pre-treated with glycolytic inhibitor 2-deoxy-D-glucose (2-DG) for 30mins (E) Cells were treated with insulin for 30mins followed by glucose for 30 min to assess induction of adipsin protein expression. (F) Experimental design of mice: 6h fasting followed by insulin (0.5IU/kg) treatment for 30 min. Plasma glucose from tail vein blood in control and insulin treated mice. (G) Immunoblots of adipsin, phosphorylated eIF2α (peIF2α), total eIF2α, phosphorylated 4EBP1 (p4EBP1), total 4EBP1, pS6, phosphorylated AKT (pAKT) and total AKT in mice liver lysates. (H) Immunoblots of peIF2α, total eIF2α from vehicle, STZ and STZ + Insulin-treated mice liver lysates. Mice n=6 (control), n=6 (insulin). Data are represented as mean±SD. *\*P*< 0.05, *\*\*P*< 0.01, *\*\*\*P*< 0.001, statistical significance was determined using unpaired two-tailed t-test or one-way ANOVA followed by Bonferroni’s multiple-comparison test as appropriate. ns, not significant.

### 2.4 Structured 5′UTR mediates mTOR-dependent, glucose-responsive translation of hepatic adipsin

Given the central role of mTOR in nutrient sensing [26; 27], we hypothesised that it might be involved in the regulation of hepatic adipsin protein synthesis. Pharmacological inhibition of mTOR by torin 1 attenuated the glucose induced hepatic adipsin protein expression, with unaltered mRNA levels (Figure 4A; Supplementary Figure 2E,F). Treatment of rapamycin, a selective mTORC1 inhibitor, also suppressed glucose induced adipsin expression (Figure 4B). Interestingly, glucose stimulation suppressed eIF2α phosphorylation even in presence of torin 1 and rapamycin and blockade of the mTOR arm of translational regulation was sufficient to inhibit adipsin expression (Figure 4A, B). Glucose mediated induction of adipsin was further augmented by the concurrent activation of mTORC1 with amino acid leucine (Figure 4C). Consistently, silencing the key mTOR complex components by siRNA against Raptor and Rictor blunted the glucose-mediated induction of adipsin protein levels (Figure 4D). Taken together, our results strongly indicate that mTOR mediates the glucose-dependent protein expression in hepatocytes.

**Figure 4.**
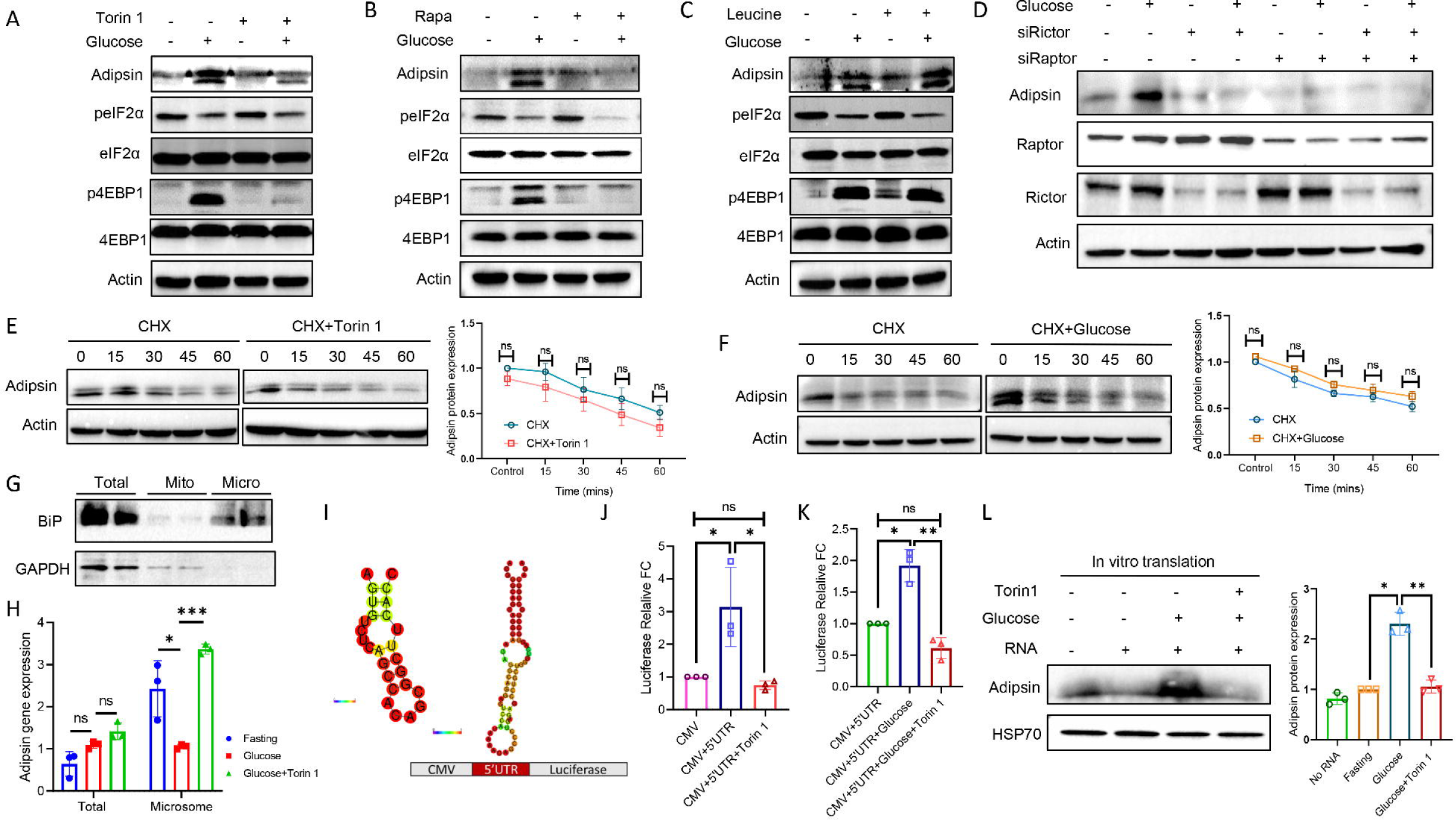
mTOR promotes adipsin translation through its structured 5′UTR in a glucose-dependent manner. (A-C) HepG2 cells were starved in HBSS for 6h prior to treatments. Cells were pre-treated with mTOR inhibitor Torin 1 (A), rapamycin (Rapa) (B), mTOR activator Leucine (C) for 30 min, followed by addition of glucose for the next 30mins to access the induction of adipsin protein expression. (D) siRNA-mediated knockdown of mTOR complex components (Raptor and Rictor) on glucose-induced adipsin protein expression. (E-F) Immunoblot analysis of HepG2 cells for adipsin protein stability assessed by cycloheximide (CHX) chase assay in the presence or absence of torin 1 (E), and glucose (F). (G) Mitochondrial and microsomal fractions from HepG2 cell lysates were isolated by differential centrifugation. Purity of microsome isolation determined by ER resident BiP by immunoblotting. (H) Quantification of adipsin mRNA expressions in total cellular and microsomal fraction under glucose deprivation, glucose stimulation, and glucose stimulation with torin 1. (I) Predicted secondary structures of murine and human 5′UTR of adipsin mRNA, and schematic of the firefly luciferase reporter construct containing the CMV promoter. (J) HEK293T cells were transfected with luciferase reporter constructs with and without 5′UTR of adipsin mRNA. After 48h, cells were treated with torin 1 and cellular luciferase activity was determined. (K) Luciferase activity in response to glucose stimulation with and without torin 1 treatment. (L) Immunoblot from a cell-free translation assay showing adipsin protein synthesis in the presence or absence of glucose and torin 1. Data are presented as mean ± SD. *\*P*< 0.05, *\*\*P*< 0.01, *\*\*\*P*< 0.001, statistical significance was determined using unpaired two-tailed t-test or one-way ANOVA followed by Bonferroni’s multiple-comparison test, as appropriate. ns, not significant; CMV, cytomegalovirus.

To investigate the mechanisms by which mTOR signaling enhances adipsin protein expression, we first assessed the stability of the adipsin protein following mTOR inhibition. Cycloheximide (CHX) chase experiments revealed that neither torin 1 nor glucose treatment significantly altered adipsin protein stability (Figure 4E, F), indicating that glucose regulates adipsin levels through a distinct mechanism. Given that secretory proteins such as adipsin are synthesized on the endoplasmic reticulum (ER)-associated ribosome for subsequent trafficking via Golgi networks [28; 29], we hypothesized that glucose may influence adipsin translation by modulatingthe association of itsmRNA with ER-bound ribosomes. To test this, we isolated cellular microsomal fractions and quantified adipsin mRNA levels (Figure 6G). Glucose deprivation led to an increased enrichment of adipsin mRNA in the microsomal fraction, an effect reversed by glucose repletion. Notably, this glucose-dependent reduction in microsomal adipsin mRNA was blocked by both torin 1 (Figure 6H), indicating that inhibition of mTOR stalls adipsin mRNA in the microsomal fraction, rendering a slower rate of translation. mTOR signaling is known to promote translation of mRNAs with structured 5′ untranslated regions (5′UTRs) by regulating the translation initiation machinery [30]. Consistent with this, RNA secondary structure prediction revealed that the 5′UTR of both murine and human adipsin mRNAs forms a stable stem-loop structure (ΔG = –25 kcal/mol, and ΔG = −5.74 kcal/mol) [31] (Figure 4I). To directly assess the functional role of the adipsin 5′UTR in translation, we generated a luciferase reporter construct containing the adipsin 5′UTR upstream of the firefly luciferase coding sequence (Figure 4I). Expression of this construct led to increased luciferase activity in an mTOR-sensitive manner (Figure 4J). Furthermore, glucose treatment enhanced luciferase activity, whereas torin 1 treatment attenuated the glucose-induced response (Figure 4K), supporting the notion that mTOR activation facilitates adipsin translation via its 5′UTR. Finally, we validated these findings in a cell-free translation system. Consistent with cellular data, glucose addition enhanced in vitro translation of adipsin, whereas torin 1 suppressed this effect (Figure 4L). Together, these results indicate that mTOR promotes adipsin protein synthesis by enhancing translation of its structured 5′UTR in a glucose-dependent manner.

### 2.5 mTOR inhibition attenuates feeding-induced hepatic adipsin protein expression

We next sought to determine whether in vivo inhibition of mTOR affects hepatic adipsin expression during the fasting refeeding cycle. To that end, mice were fasted for 12 h and subsequently administered with torin 1(10mg/kg body weight, intraperitoneal) for 4 h [32]. A subset of mice was refed during the final hour of this period (Figure 5A). As expected, torin 1-treated animals exhibited elevated fasting blood glucose levels compared to controls in both fasted and refed conditions, accompanied by a concurrent increase in circulating insulin levels (Figure 5B, C). Western blot analysis revealed that hepatic adipsin protein levels were markedly reduced in both fasted and refed groups following torin 1 treatment, paralleling decreased phosphorylation of p4EBP1, a downstream target of mTORC1(Figure 5D). Adipsin mRNA levels were expected to remain unaltered (Figure 5E). Immunohistochemical analysis further confirmed the reduction in hepatic adipsin protein expression(Figure 5F). We also assessed circulating adipsin levels and found a significant decrease in serum adipsin concentrations in torin1-treated mice, suggesting that mTOR inhibition suppresses adipsin production in both hepatic and adipose tissues (Figure 5G). Importantly, no significant differences were observed in body weight, liver weight, or adipose tissue mass across treatment groups (Figure 5H, I). Collectively, these findings demonstrate that hepatic adipsin protein expression is positively regulated by mTOR signaling during the postprandial phase, primarily through translational mechanisms rather than transcriptional changes.

**Figure 5.**
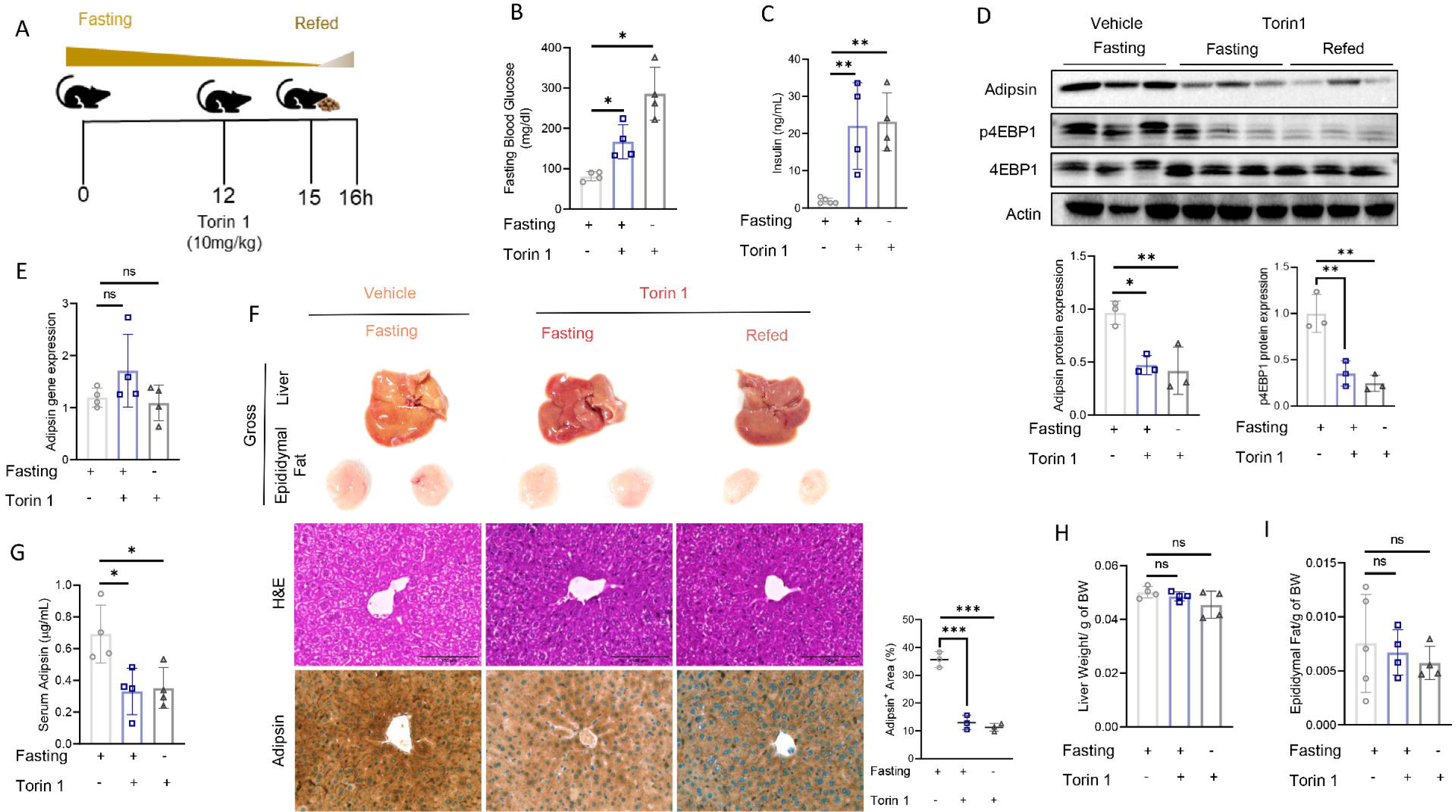
Pharmacological inhibition of mTOR suppresses hepatic adipsin expression during the fasting–refeeding cycle. (A) Experimental design depicting 12 h fasting followed by 4 h treatment with torin 1 (10 mg/kg, intraperitoneal), with a subset of mice refed during the final hour. (B–C) Plasma glucose and insulin concentrations in vehicle- and torin 1-treated mice under fasted and refed conditions. (D) Immunoblot analysis of hepatic adipsin and phosphorylated S6 (pS6) protein levels in response to torin1 treatment. (E) Quantification of hepatic adipsin mRNA expression following torin 1 treatment. (F) Gross liver and epididymal adipose tissue, and liver sections for H&E, immunohistochemistry of adipsin. Adipsin positive area (%) was quantified (n=3/group, 6 fields/sample). (G) Serum adipsin concentrations in vehicle and torin 1-treated mice. (H–I) Body weight, liver weight, and epididymal adipose tissue depot across experimental groups. Mice n=6(vehicle), n=6(torin 1, fasting), n=6 (torin1, refed). Data are presented as mean ± SD. *\*P*< 0.05, *\*\*P*< 0.01, *\*\*\*P*< 0.001, statistical significance was determined using one-way ANOVA followed by Bonferroni’s multiple-comparison test. Scale bar, 100 μm. ns, not significant.

### 2.6 Depletion of adipsin in the mouse liver promotes glucose intolerance

To investigate the physiological role of hepatic adipsin in metabolic homeostasis, we selectively silenced its expression in the liver using adenovirus-mediated shRNA (Supplementary Figure 3A, B). As expected, adipsin mRNA expression (Figure 6A) was significantly lower in the adenovirus-mediated shAdipsin group compared to the shGFP. Consistently, histological and immunohistochemical analyses showed a significant downregulation in adipsin levels (Supplementary Figure 3C). We first examined whether hepatic adipsin depletion affects basal metabolic parameters. Interestingly, hepatic adipsin knockdown did not alter fasting blood glucose levels and circulating insulin concentrations (Figure 6B, C). However, glucose tolerance test (GTT) revealed a markedly reduced glucose clearance in the shAdipsin group, as reflected by a higher area under the curve (AUC) compared with shGFP-treated mice (Figure 6D).To determine whether this impaired glucose tolerance is due to defects in insulin secretion, we performed glucose-stimulated insulin secretion (GSIS). Despite the known role of adipsin in improving β-cell function (3, 11), insulin secretion in response to glucose challenge was not altered in the shAdipsin group (Figure 6E). We next assessed insulin responsiveness, and found that insulin-dependent glucose lowering and downstream signaling remained unchanged (Figure 6F, Supplementary Figure 3D). We further evaluated whether changes in body composition or lipid metabolism could explain the observed phenotype. Body weight, liver weight, epididymal adipose tissue, and total triglyceride levels were comparable between groups (Supplementary Figure 3E–H), indicating that the impaired glucose tolerance is not due to alterations in overall metabolic status. Since hepatic glucose production is a major determinant of systemic glucose levels [33], we next examined gluconeogenesis. We observed that expression of gluconeogenic genes and proteins remained unchanged (Supplementary Figures 3I and 3J), suggesting that hepatic glucose output is not affected by adipsin depletion. Having excluded alterations in insulin secretion, insulin sensitivity, body composition, and gluconeogenesis, we hypothesized that impaired hepatic glucose uptake may underlie the observed phenotype, and examined the hepatic glucose transporters. Expressions of glucose transporters, specifically glucose transporter type 2 (GLUT2), were significantly downregulated upon adipsin knockdown (Figure 6G, H). Consistently, in vitro glucose uptake assays showed a marked reduction in glucose uptake in both HepG2 (1.4-fold) and AML12 (4.5-fold) hepatocytes, indicating a cell-autonomous effect. Adipsin silencing in AML12 cells reduced GLUT2 mRNA and protein levels. Conversely, overexpression of adipsin in AML12 cells enhanced glucose uptake (Supplementary Figure 4A), and increased the protein levels of adipsin and GLUT2 (Supplementary Figure 4B). To further validate these findings in vivo, we overexpressed hepatic adipsin in mice (Supplementary Figure 4C). As expected, hepatic adipsin expression was markedly elevated, accompanied by increased circulating adipsin levels. Consistent with our in vitro observations, hepatic GLUT2 protein levels were also increased in adipsin-overexpressing mice (Supplementary Figure 4D, E). We next assessed metabolic parameters and found no significant changes in fasting blood glucose, circulating insulin levels, body weight, liver weight, total triglycerides, or cholesterol levels between groups (Supplementary Figure 4F-K). Interestingly, GTT, as reflected by a higher area under the curve (AUC) compared with Ad-eGFP-treated mice, revealed significantly improved glucose clearance in adipsin-overexpressing mice (Supplementary Figure 4L), supporting a functional role for hepatic adipsin in promoting hepatic glucose uptake and maintaining systemic glucose homeostasis.

**Figure 6.**
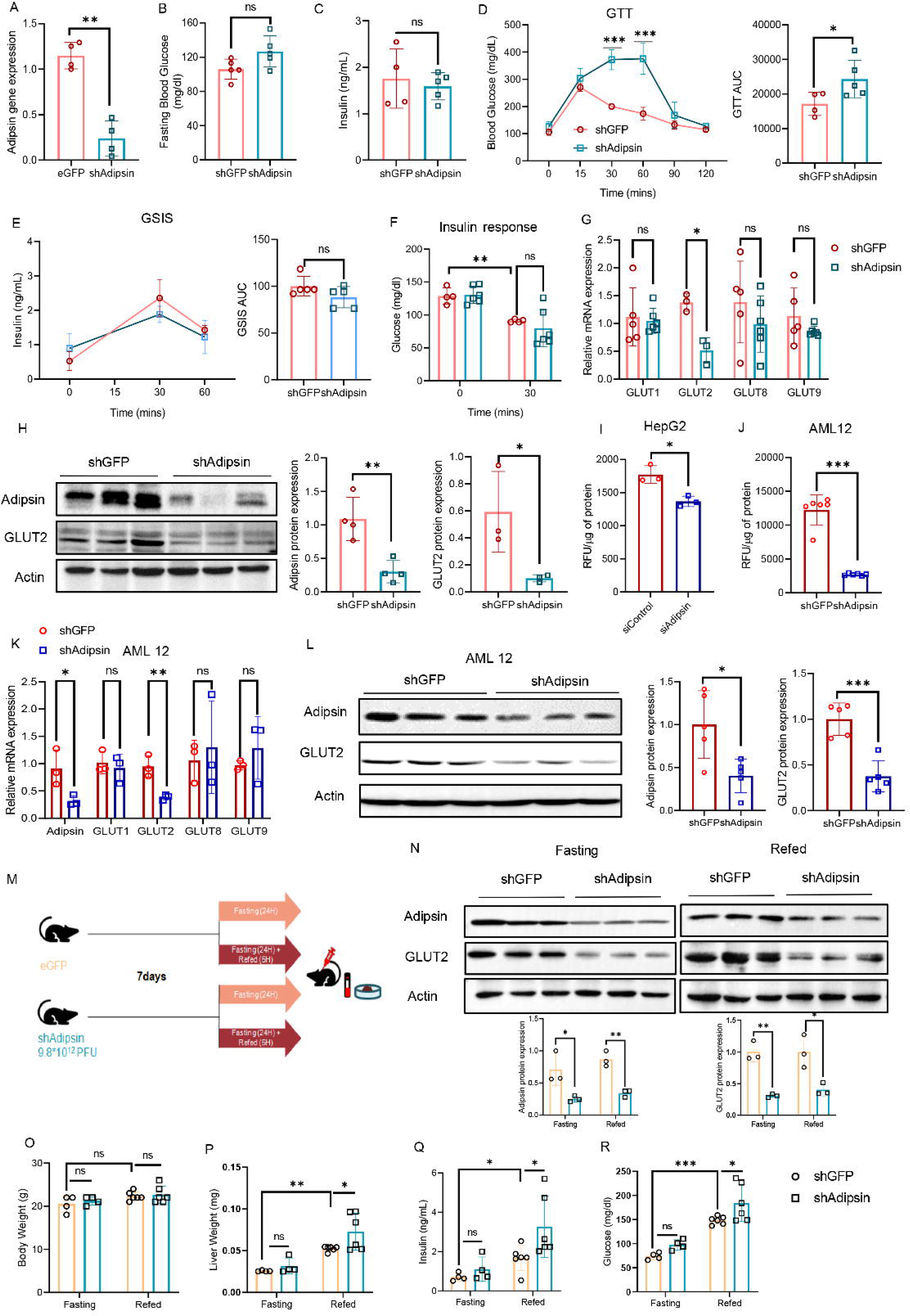
Hepatic adipsin deficiency impairs systemic glucose tolerance and postprandial metabolic regulation. (A) mRNA expression of adipsin in liver homogenates following adenovirus-mediated shRNA knockdown of Adipsin. (B, C) Fasting blood glucose levels(B), plasma insulin concentrations (C)in shGFP and shAdipsin-treated mice. (D-G) Glucose tolerance test (GTT), and corresponding area under the curve (AUC) (D), Glucose-stimulated insulin secretion (GSIS)(E), insulin tolerance test (ITT) (F), mRNA expression of GLUT1, GLUT2, GLUT8, and GLUT9 genes (G) in shGFP and shAdipsin-treated mice. (H) Immunoblot analysis of adipsin and GLUT2 protein in hepatic homogenates following adenovirus-mediated shRNA knockdown of Adipsin. Right panel: Densitometric analysis of Adipsin and GLUT2. (I) Glucose uptake assay in HepG2 cells following siRNA-mediated knockdown of adipsin. (J) Glucose uptake assay in AML12 cells following adenovirus-mediated shRNA knockdown of Adipsin. (K) mRNA expression of Adipsin, GLUT1, GLUT2, GLUT8, and GLUT9 genes in shGFP and shAdipsin-treated AML12 cells. (L) Immunoblot analysis of adipsin and GLUT2 protein in hepatic homogenates following adenovirus-mediated shRNA knockdown of Adipsin in AML12 cells. Right panel: Densitometric analysis of Adipsin and GLUT2 (n=6). (M) Experimental design for the fasting–refeeding protocol. (N) Immunoblot analysis of hepatic adipsin and GLUT2 expression following fasting and refeeding. Right panel: Densitometric analysis of adipsin and GLUT2 expression (n=3). (O-R) Quantification of body weight (O), liver weight (P), plasma insulin (Q) and plasma glucose concentrations (R) in shGFP and shAdipsin-treated mice subjected to fasting–refeeding. Mice n = 10 (shGFP), n = 10 (shAdipsin), n= 10 (fasting, shGFP), n= 10 (fasting, shAdipsin), n= 10 (refed, shGFP), n= 10 (refed, shAdipsin). Data are presented as mean ± SD. *\*P*< 0.05, *\*\*P*< 0.01, *\*\*\*P*< 0.001, statistical significance was determined using unpaired two-tailed t-test or one-way ANOVA followed by Bonferroni’s multiple-comparison test, as appropriate. ns, not significant; shGFP, Green Fluorescent Protein.

**Figure 7.**
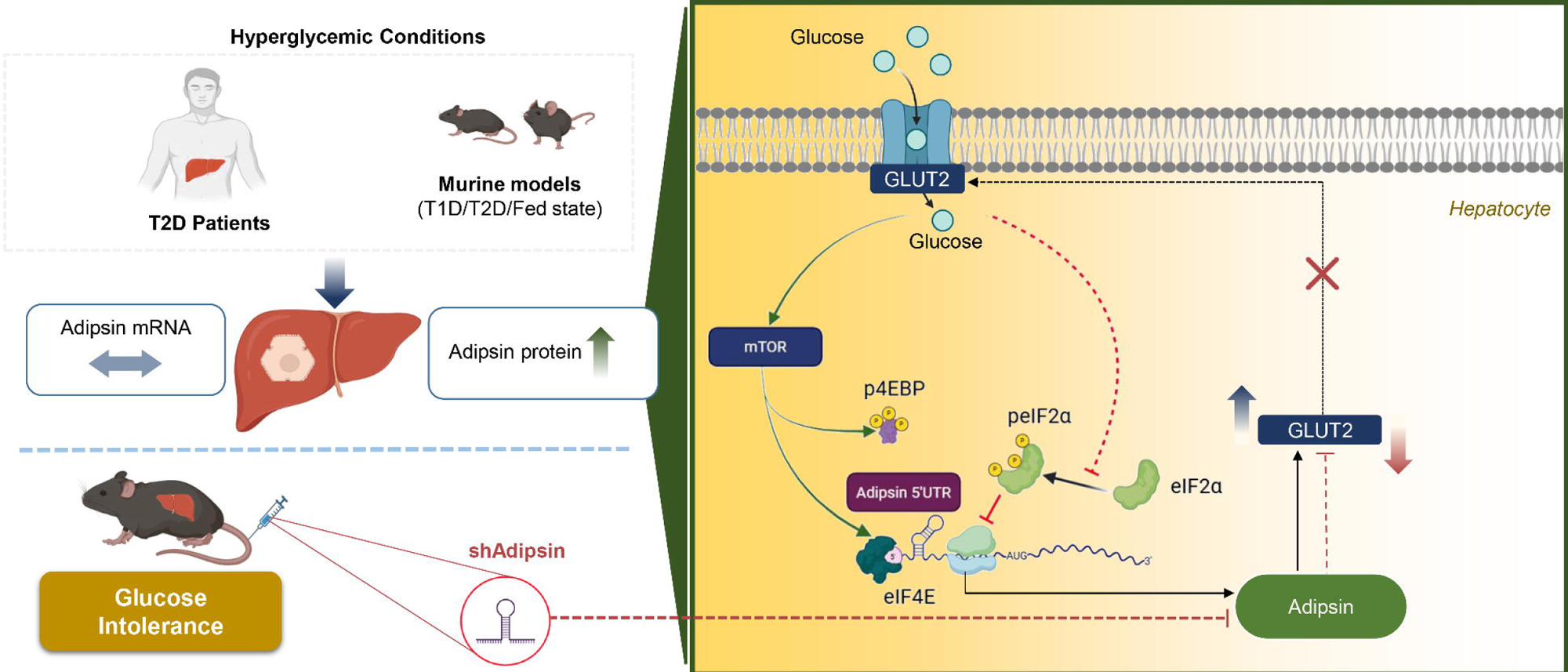
Schematic of glucose dependent hepatic adipsin expression, and its implication in diabetes and glucose metabolism.

Next, we examined whether adipsin knockdown alters glucose excursions during the fasting–refeeding cycle. Mice with hepatic adipsin silencing were subjected to a fasting–refeeding protocol (Figure 6M). As expected, adipsin and GLUT2 protein levels were significantly reduced in the adipsin knockdown group (Figure 6N). Interestingly, adipsin silencing increased adipose tissue mass (Supplementary Figures 4M, N), whereas body weight and liver weight (Figures 6O, P), as well as circulating triglyceride and cholesterol levels, remained unaltered between groups (Supplementary Figure 4O, P), thus suggesting that hepatic adipsin depletion does not markedly disrupt systemic lipid metabolism responses. Refeeding increased circulating insulin and glucose levels in adipsin-deficient mice (Figures 6Q, R). In conjunction with impaired glucose tolerance during GTT and reduced GLUT2 expression and hepatocyte glucose uptake, these findings indicate that hepatic adipsin depletion disrupts postprandial glucose handling and impairs systemic glucose tolerance. Collectively, our data support a model in which the glucose–mTOR axis governs hepatic adipsin translation, while hepatic adipsin, in turn, facilitates efficient glucose uptake and systemic glycemic control, forming a reciprocal regulatory loop.

## 3. Discussion

The immunometabolic functions of adipsin have been implicated in multiple metabolic disorders [34–37], although prior studies have yielded inconsistent results [20; 21; 38-40]. These discrepancies underscore the necessity of delineating its tissue-specific expression patterns and regulatory mechanisms. In this study, we uncover previously unrecognized aspects of hepatic adipsin biology in the context of glucose homeostasis. We demonstrate that both human and murine livers serve as significant sources of adipsin, with protein levels rapidly induced in the postprandial state and in diabetes, despite unchanged mRNA abundance. Furthermore, we identify a mechanistic link whereby glucose stimulates hepatic adipsin translation through coordinated regulation of eIF2α and mTOR signaling, with the 5′UTR of adipsin mRNA conferring mTOR-dependent translational control. Importantly, hepatocyte-specific depletion of adipsin impaired systemic glucose tolerance, particularly in the postprandial state. This defect was associated with reduced hepatic glucose uptake, accompanied by a marked downregulation of GLUT2. Collectively, these findings establish hepatic adipsin as a glucose-responsive regulator that links nutrient sensing to hepatic glucose handling.

Our data further demonstrate that liver-derived adipsin is not merely a by-product of metabolic stress but functions as an active regulator of systemic glucose homeostasis [11]. Genetic ablation of hepatic adipsin resulted in impaired glucose tolerance without alterations in insulin secretion or signaling, indicating that the observed phenotype is not driven by defects in β-cell function or insulin responsiveness. Instead, the reduction in GLUT2 expression and hepatocyte glucose uptake suggests a primary defect in hepatic glucose handling. While adipose-derived adipsin has largely been considered a systemic endocrine factor that enhances β-cell function via complement C3a–mediated mitochondrial support[3; 11; 41; 42], our findings extend this paradigm by identifying a tissue-specific role for hepatic adipsin in facilitating glucose uptake. These results reconcile previously conflicting reports on circulating adipsin and metabolic liver disease [39] by demonstrating that hepatic adipsin is subject to post-transcriptional regulation and exerts local metabolic effects that may not be reflected by circulating levels. Such context-dependent regulation may explain reports associating elevated adipsin levels with increased MASLD severity [18], as well as studies describing inverse relationships [21]. Together, these observations suggest that circulating adipsin concentrations may not reliably reflect its tissue-specific function and that its role may vary across disease stages, potentially acting as a compensatory factor during early metabolic stress but becoming dysregulated in chronic disease [43; 44].The precise consequences of hepatocyte-derived adipsin on the liver microenvironment and overall hepatic physiology, however, remain to be fully elucidated.

Mechanistically, our data indicate that glucose, through its metabolism, simultaneously impacts eIF2α and mTOR, both of which are required for effective adipsin synthesis. Glucose deprivation rapidly induces eIF2α phosphorylation [24], leading to inactivation of the eIF2 trimer, and its interaction with the initiator tRNA and engagement with the start codon[45]. Concurrently, inhibited mTOR complex hinders uncoupling of eIF4E from its binding partner 4E-BP1, and binding to 5’ cap of adipsin mRNA, thereby suppressing translation initiation[46].Of these two controlling arms, the glucose–mTOR signalling axis promotes adipsin protein translation via the 5′UTR of adipsin mRNA. Potential mechanisms may involve secondary RNA structures or internal ribosome entry sites (IRES) within the 5′UTR that integrate nutrient-sensitive signaling cascades, as has been described for other mTOR-dependent transcripts [47; 48].Functionally, our findings place hepatic adipsin at a critical node linking nutrient sensing to glucose metabolism. In the context of preserved insulin secretion and signaling, the observed impairment in glucose tolerance following adipsin depletion can be attributed to defective hepatic glucose uptake, highlighting a previously unappreciated role for adipsin in regulating GLUT2 expression and glucose uptake. Ribosome profiling and reporter-based assays will be instrumental in identifying such elements and in clarifying how nutrient signaling intersects with adipsin synthesis. Furthermore, the downstream metabolic functions of adipsin—as both an endocrine factor and a tissue-specific modulator—remain incompletely understood. Future studies should address the relative contributions of alternative complement activation versus direct modulation of hepatocellular pathways, including glucose transport, in mediating the metabolic effects of adipsin.

In conclusion, this study identifies liver-derived adipsin as a nutrient-responsive protein whose translation is governed by the glucose–eIF2α–mTOR axis and that plays a direct functional role in systemic glucose homeostasis. By linking glucose-dependent translational regulation of adipsin to hepatic glucose uptake and GLUT2 expression, our findings provide a mechanistic basis for its contribution to glucose intolerance. These results establish the liver as a key source of adipsin regulated through post-transcriptional mechanisms and expand the current understanding of its role beyond adipose tissue. Collectively, our work positions hepatic adipsin as a tissue-specific regulator of glucose metabolism with potential therapeutic relevance in metabolic liver disease and diabetes.

## Author contributions

SKM: Methodology, experimentation, data analysis, visualization, writing the originaldraft; AS: Generation of plasmid construct and experimentation. TD, AB, and AS: biochemical and in vivo experiments. SM and AC: patient recruitment, clinical classification, liver biopsy, and analysis of histology. PC: Conceptualization, designed study, researched data, methodology, writing & editing, project administration, funding acquisition; PC is the guarantor of this work and, as such, had full access to all the data in the study and takes responsibility for the integrity of the data and the accuracy of the data analysis.

## Acknowledgments

The authors thank the Central Instrumentation Facility (CIF) of CSIR-IICB, Rabin Pramanik, for maintaining animal cohorts and Tapan Das for assistance with histology. SKM thanks ICMR, India, for his research fellowship. The authors have declared that no conflict of interest exists.

## Data Availability

All study data are included in the manuscript and/or supporting information.

## Materials and Methods

### Human subjects

A total of 30biopsy-proven patients diagnosed with MASLD were recruited for this study. Following participants’ written consent, liver biopsies were collected from patients in the Department of Gastroenterology at the Indian Institute of Liver and Digestive Sciences (IILDS), West Bengal, India (IEC reference No. HREC, IILDS/2023-R11). Study participants’ inclusion criteria were with or without diabetes, and exclusion criteria included: any liver disease other than MASLD (i.e., alcoholic liver disease, hepatitis B or C, autoimmune hepatitis, hemochromatosis, Wilson disease, or drug-induced hepatitis). Histopathological parameters, including steatosis score, hepatocellular ballooning, fibrosis stage, and NAS, were determined by a single pathologist. Fasting blood samples were collected from the patient for HbA1c, and plasma glucose, insulin, total triglycerides, cholesterol, AST, and ALT were measured. HOMA-IR was calculated as (fasting insulin × fasting glucose)/22.5; and HOMA-β was calculated as (20 × fasting insulin)/ (fasting glucose − 3.5). Plasma and liver adipsin levels were detected by an ELISA kit (R&D Systems) according to the manufacturer’s protocol.

### Animal Experiments

Six- to eight-week-old C57BL/6 male mice were kept in ventilated cages at ambient temperature with a 12:12-hour light-dark cycle at 22±1°C, with unrestrained access to food and water. All experiments and protocols were carried out in accordance with the Institutional Animal Ethics Committee at the CSIR-IICB (approved by the CPCSEA, Ministry of Environment & Forest, and Government of India). After acclimatization, for each experiment, mice were weighed and then randomized accordingly to ensure similar mean body weight between groups. For the Streptozotocin (STZ) experiment, mice were treated with a single intraperitoneal dose of STZ (150mg/kg of body weight). After the fifth day, subcutaneous insulin (0.75U/kg, Actrapid, Novo Nordisk Pharmaceuticals Ltd., Auckland) was administered twice daily for the next seven days to achieve glycaemic control. Mice were fed ad libitum with 60%high fat diet (HFD; D12492, Research Diets, Inc., 20 Jules Lane, New Brunswick, NJ 08901 USA) or a chow diet for 16 weeks. A 24h fasting in mice was performed with total food deprivation and ad libitum access to water from 1200h to 1200h of the following day, following which one group of mice was refed for 5 h and then sacrificed. For the insulin experiment, mice were fasted for 6h and then randomly divided into two groups. The insulin-treated groups received 0.75 IU/kg of body weight of insulin and were sacrificed after 30 min.

### Adenovirus

For liver-specific knockdown of Adipsin, six to eight-week-old male mice (C57BL/6) were intravenously injected with purified shRNA construct or shGFP at a titer of 9.8*10^12^ pfu/mouse, generated by BLOCK-iT Adenoviral Expression System (Invitrogen) using the sequence #5’-CACCGCCTGATGTCCTGCATCAACTCGAAAGTTGATGCAGGACATCAGGC-3’. Mouse CFD-HA adenovirus was purchased from ABM (# 159690540200) and used for overexpression of adipsin in cells and mouse liver.

### Glucose tolerance test (GTT)and glucose-stimulated insulin secretion (GSIS)

GTT was performed as described by the Mouse Metabolic Phenotyping Centers (MMPC). Briefly, mice were fasted for 6 h with free access to water, and fasting blood glucose levels weremeasured using a glucometer (SD CodeFree). Mice were then injected with 2 g/kg of glucose, and blood glucose levels were measured at 15, 30, 60, and 120 minutes after glucose injection. For GSIS, insulin was measured at 0, 30, and 60 minutes.

### Cell Culture

Primary hepatocytes were isolated from 6-8-week-old C57BL/6 mice as described [49].Briefly, the liver was perfused with HBSS without calcium, followed by a pre-warmed solution of type I Collagenase (1 mg/mL, Roche) in HBSSwith calcium. Cell suspension was centrifuged at 50 g for 2 min at 4°C to pellet down the parenchymal cells, and seeded onto a collagen-coated tissue culture plate. Human hepatoma cell line HepG2 was cultured in Minimum Essential Medium (HiMedia, India) supplemented with 10% Fetal Bovine Serum (Gibco, USA) and 1% Penicillin-Streptomycin. Cells were treated with L-Leucine (L8000, Sigma Aldrich, USA), Insulin Human Recombinant (#91077C, Sigma Aldrich, USA), Torin 1 (#4247, Tocris Biosciences, USA), Rapamycin (#13346, Cayman chemicals, USA), and 2-DG (#14325, Cayman chemicals).

### Microsome Isolation

HepG2 cells were scraped into ice-cold PBS and centrifuged at 600 g for 5 min at 4 °C. The cells were then resuspended in hypotonic buffer (10 mM HEPES, 1 mM EGTA, 25 mMKCl) and centrifuged at 600 g at 4 °C for 5 min. The cells were resuspended in isotonic buffer (10 mM HEPES, 1 mM EGTA, 25 mMKCl, 250 mM sucrose) and homogenized. Following which centrifugation was performed at 1000g, 4 °C for 10 min. The supernatant was transferred to a new tube and then centrifuged at 12000g, 4 °C for 15 min to collect the mitochondrial fraction. The supernatant was then transferred, 7.5 times CaCl2 (8 mM) was added, and the sample was placed in a slow rotor for 45 min. Centrifugation was then performed at 8000 g at4 °C for 10 min to collect the microsomes. RNA isolation was then performed from the microsomes.

### Quantitative RT-PCR

Total cellular and tissue RNA was isolated using TRIzol reagent (Invitrogen). TheiScriptc DNA Synthesis Kit prepared cDNA, and gene expression was quantified by using SYBR green mix (SYBR® Green supermix Bio-Rad) in the Light Cycler 96 real-time PCR System (Roche, Switzerland). The sequences of primers used-Human Adipsin: Forward 5’-CTACAGCTGTCGGAGAAG-3, Reverse 5-CCGCGTGGTTGACTATG-3’; Mouse Adipsin: Forward 5’-GGTATGATGTGCAGAGTGTAG-3’, Reverse: 5’-GGTTCCACTTCTTTGTCCTC-3’; Mouse PCK1: Forward 5’-GAAGAGGACTTTGAGAA-3’,Reverse: 5’-GTCAGTTCAATACCAATC-3’; Mouse Creb1: Forward 5’-CTCCCACTGTAACCTTAGT-3’, Reverse: 5’-CTGTTTGGACTTGTGGAG-3’; Mouse Fbp1: Forward 5’-CAAAGCCATCTCGTCTG-3’, Reverse: 5’-GTATGTCCAGCTTCTTTACT-3’; Mouse Fbp2: 5’-GAATGTGACAGGAGATGAG-3’, Reverse 5’-CACCGCCTCTTTATTCTC-3’; Mouse G6pc1: Forward 5’-CTGTGGGCATCAATCTC-3’. Reverse 5’-GCTGTTGCTGTAGTAGTC-3’; Mouse G6pc3: Forward 5’-GGCTCAACCTTGTCTTC-3’, Reverse 5’-AGGGAACTGGTGAATCT-3’; Mouse Mdh1: Forward 5’-ACGGACAAAGAAGAGATTG-3’, Reverse 5’-TCACATTGGCTTTCAGTAG-3’; Mouse Mdh2: Forward 5’-CCAGATTGCCTCAAAGG-3’, Reverse 5’-GTAGCGTTGGTGTTGAA-3’; Mouse Pck1: Forward 5’-CGCAAGCTGAAGAAATATG-3’, Reverse 5’-GCTCTTGGGTGATGATG-3’; Mouse Pck2: Forward 5’-CTGAGAACACTGCCATAC-3’, Reverse: 5’-ATCACCGTCTTGCTTTC-3’; Mouse Pfkfb1: Forward 5’-GTGAGCTACAGGAACTATG-3’, Reverse 5’-CACGGCTGAGATACTTATG-3’; Mouse Ppara: Forward 5’-ACCTGGAAAGTCCCTTAT-3’, Reverse 5’-TCCTAAGTACTGGTAGTCTG-3’; Human RN 18S: Forward 5’-GTAACCCGTTGAACCCCATT-3’, Reverse 5’-CCATCCAATCGGTAGTAGCG-3’. Mouse RN 18S: Forward 5’-GTTGGTTTTCGGAACTGAGG-3’, Reverse: 5’-GTAGCCACCACCTCCCAGTA-3’

### Immunoblotting

Cellular and liver protein extracts in lysis buffer [50 mM Tris-HCl (pH 7.4), 100 mM NaCl, 1 mM ethylenediaminetetraacetic acid (EDTA), 1 mM ethylene glycol-bis(β-aminoethyl ether)-N,N,N′,N’-tetraacetic acid (EGTA), and 1% Triton X-100 along with protease- and phosphatase inhibitor cocktail (Roche, Basel, Switzerland)] was resolved using SDS-PAGE followed by transfer to PVDF Membrane (Immobilon-P, Merck Life Sciences, NJ, USA). Membranes were incubated overnight with primary antibodies, followed by incubation with peroxidase-conjugated secondary antibodies, and the bands were visualized with a ChemiDoc MP Imaging System (Bio-Rad, Hercules, CA, USA). Following primary antibodies were used: Human Adipsin (#A21110), Mouse Adipsin (#A8117), PCK1 (#A22172) was purchased from ABclonal (MA, USA), pS6Ribosomal Protein (#4858), S6Ribosomal Protein (#2217), Raptor (#2280), Rictor (#2114), pAKT(#4060), AKT(#4691), peIF2α (#3398), eIF2α (#5324),p4E-BP1 (#2855), 4E-BP1 (#9644), BiP (#3183), GAPDH (#2118)were purchased from Cell Signalling Technology (Minneapolis, USA). β-actin (#A2228) was purchased from Sigma-Aldrich (St. Louis, Missouri, United States).

### Histology, Immunohistochemistry (IHC), Periodic Acid-Schiff stain (PAS)

The liver tissue was excised immediately and immersed in 10% formalin. The tissue blocks were then sectioned using a rotary microtome with a thickness of 5µm. Deparaffinization was performed by baking the tissue section at 80°C for 20 min and then rehydrating and processing for hematoxylin & eosin (H&E), and PAS, while for IHC it was followed by antigen retrieval in Tri-sodium citrate buffer (10 mM sodium citrate, 0.05% Tween 20, pH 6.0). The slides were incubated with the primary antibody, then with a diluted biotinylated secondary antibody, and stained with anti-rabbitVECTASTAIN ABC KIT.

### 5’UTR construct, Transfection, and Luciferase assay

Secondary structures of human and mouse adipsin 5’UTR were predicted by using the RNAfold webserver (31).To obtain Firefly Luciferase (FL) expression from a CMV promoter, the CMV promoter region was amplified by using *Kpn*I-*Nhe*I sites (*Kpn*I: 5’-GGGGTACCGACATTGATTATTGACTAGT-3’; *Nhe*I: 5’-CGGCTAGCAGCTCTGCTTATATAGACCT-3’) from pcDNA3.1 myc-His B plasmid. It was then inserted into the pGL3-Basic plasmid to get pGL3-CMV-FL. To generate the 5’UTR of adipsin mRNA, cDNA from mice primary hepatocytes was amplified (pre-incubation: 95°C 10min, denaturation at 95°C for 10s, annealing at 64.5°C for 30s, extension at 72°C for 45s for 30cycles) using *Nhe*I-*Hind*III sites (*Nhe*I: 5’-CTAGCTAGCAGGGGCAGGAGGTAAG AG-3’ and *Hind*III: 5’-CCCAAGCTTTCTGACAGCAGGCACTG-3’). The amplified product was then inserted downstream of CMV in pGL3-FL-CMV to generate pGL3-CMV-FL-5’UTR.Constructs were transfected in HEK293T cells using Lipofectamine^TM^ 2000 Transfection Reagent (Thermo Fisher Scientific, MA, USA) for 48 h. Luciferase activity was detected with the Dual-Luciferase Reporter Assay System (Promega) according tothe manufacturer’s protocol.

### Glucose uptake assay

Glucose uptake assay was performed by treating the HepG2 or AML12 cells with 100 µM of 2-NBDG (2-(N-(7-nitrobenz-2-oxa-1,3-diazol-4-yl) amino)-2-deoxyglucose) (Invitrogen) for 20 min. The cells were then lysed, and relative fluorescence was measured using a fluorimeter (BioTek Synergy H1 Multimode Reader, Agilent) at an excitation wavelength of 475 nm and an emission wavelength of 550 nm.

### Cycloheximide (CHX) Chase assay

The HepG2 cells were treated with either 50 μg/ml of cycloheximide (Calbiochem, Merck Millipore, MA, USA) alone or in combination with torin 1 and glucose for the required time periods. Cells were then harvested in protein lysis buffer at different time points for Western blotting.

### In vitro translation

In vitro translation assay was performed according to the manufacturer’s protocol (PROMEGA Rabbit Reticulocyte Lysate System, Nuclease Treated, L4960). Briefly, total cellular RNA was isolated and added to the reticulocyte lysate with amino acids at a final concentration of 200 µg/mL.In vitro translation was then performed by keeping the tubes at 30 °C for 90 min. Immunoblotting was performed to detect protein synthesis.

### Biochemical tests and liver enzymes

Blood samples were centrifuged at 2000g for 10 min, and plasma/serum were assayed using total triglycerides, cholesterol, glucose, AST, and ALT reagent kit (TR2774, CH2773, GL2797; AS1204, AL2780; Randox Laboratories Limited, UK) and were measured in a semi-auto analyzer (Microlab 300; ELITech Group, France). Human and mouse adipsin ELISA kit (R&D Systems, Cat: DY5430, DY1824) and insulin ELISA(Millipore Cat: EZHI, EZRMI) were performed using the manufacturer’s protocol.

### RNA interference (RNAi)

siRNA transfection was carried out using LipofectamineTMRNAiMax Transfection Reagent (Thermo Fisher Scientific, MA, USA) for 72 h. Human Raptor siRNA SMARTPOOL (Cat No.L-004107-00-0005) and Human Rictor siRNA SMARTPOOL (Cat No.L-016984-00-0005) were purchased from Dharmacon, Horizon Discovery (Waterbeach, UK).

### Statistics

Unless otherwise stated, data are presented in mean±SD. Statistical analysis was carried out using GraphPad Prism 8 and RStudio 8.0.2 (263) and RStudio Version 4.3.3. Unpaired t-test with Welch correction and ANOVA were used as appropriate, followed by Bonferroni post-hoc test. Differences were considered significant at p <0.05.

