## Supplementary Files for "Glucose-dependent regulation of hepatic adipsin controls glucose uptake and tolerance"

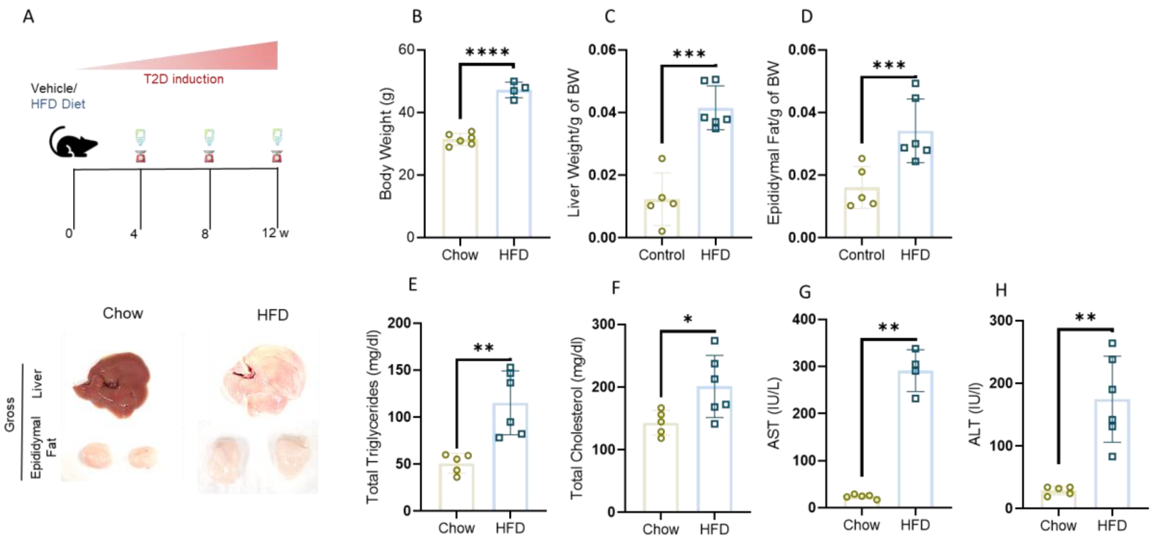


**Supplementary Figure 1. Metabolic characterization of mice following high fat diet (HFD) feeding.** (A) Top: Experimental design of HFD diet fed mice. Bottom: Gross liver and epididymal adipose tissue depot. (B-H) Body weight (B), liver weight (C), adipose tissue weight (D), serum triglycerides (E), serum cholesterol (F), AST (G), ALT (H), in Chow fed and HFD fed diet mice. Data are presented as mean ± SD.*$P< 0.05$, **$P< 0.01$, ***$P< 0.001$, statistical significance was determined using unpaired two-tailed t-test.

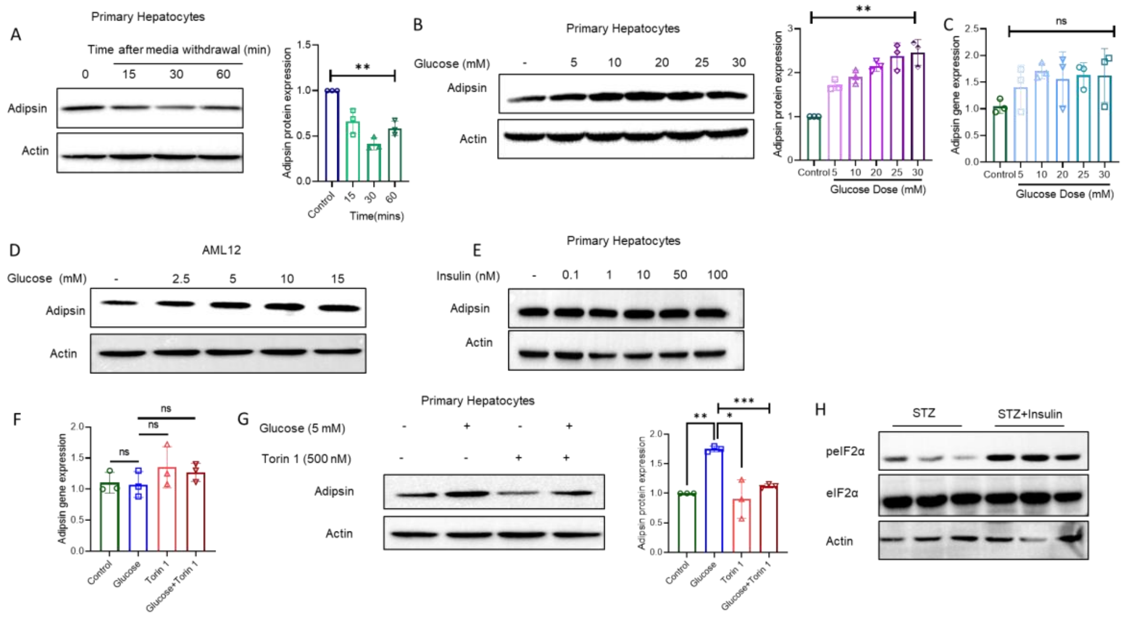


**Supplementary Figure 2. Hepatic adipsin expression is glucose-dependent and suppressed by mTOR inhibition.** (A) Immunoblot of adipsin in primary hepatocytes following media withdrawal for the indicated time points. Right: Densitometric analysis of relative adipsin protein levels. (B) Dose-dependent effects of glucose on adipsin protein expression. Right: Densitometric analysis. (C) Adipsin mRNA levels across increasing glucose concentrations. (D) Dose-dependent effects of glucose on adipsin protein expression in AML12 cells. (E) Insulin dose–response in primary hepatocytes. (F) Treatment with mTOR inhibitor Torin 1. Right: densitometric analysis of adipsin protein levels. (G) Adipsin mRNA levels following Torin 1 treatment. (H) Immunoblots of peIF2α, total eIF2α from STZ and STZ + Insulin-treated mice liver lysates. Mice n=6 (control), n=6 (insulin). Data are presented as mean ± SD.***P*< 0.01,statistical significance was determined using unpaired two-tailed t-test. ns, not significant.

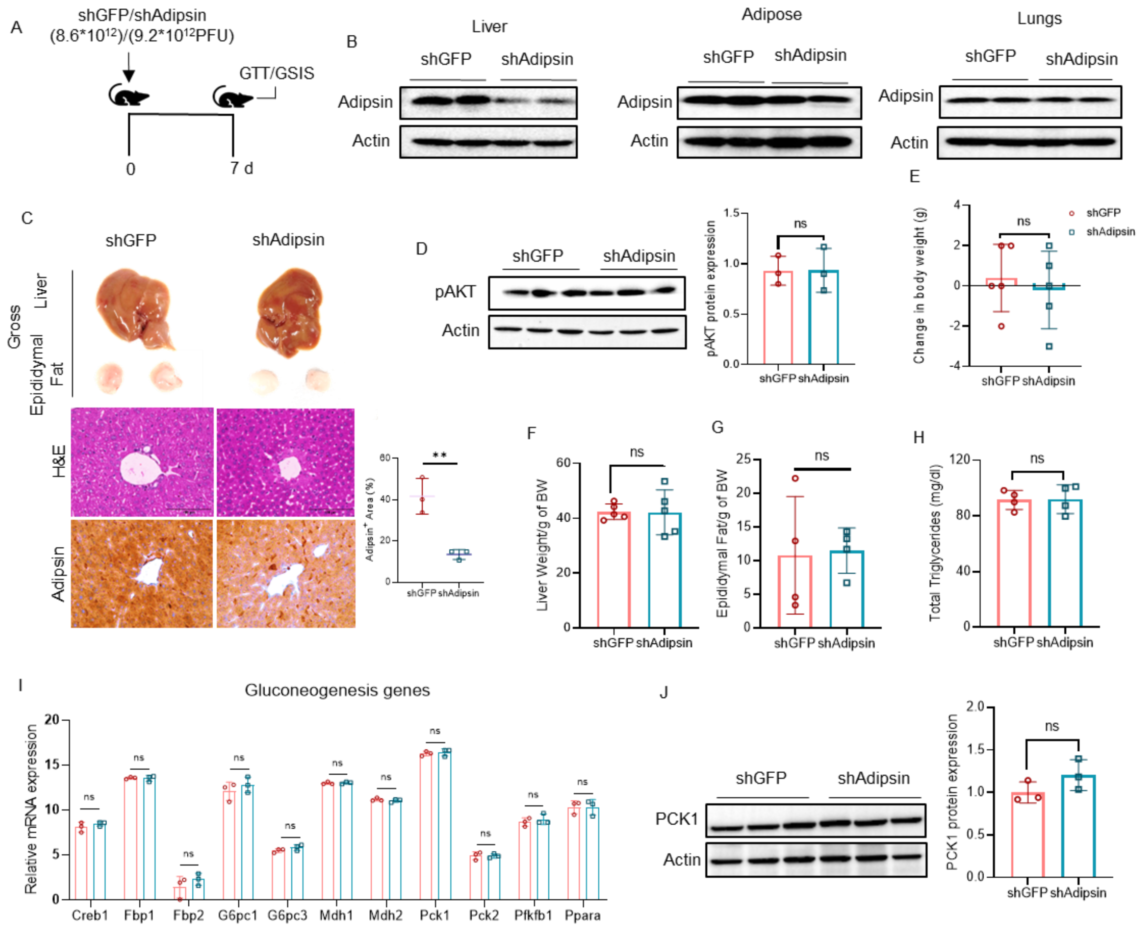


**Supplementary Figure 3. Hepatic adipsin depletion does not alter body composition or hepatic lipid content.** (A) Experimental design of adenovirus-mediated shRNA knockdown of hepatic Adipsin. (B) Adipsin knockdown via adenoviral delivery in liver, adipose, and lungs. (C) Representative gross liver and epididymal adipose tissue depot, H&E staining, and immunohistochemistry for hepatic adipsin; adipsin-positive area (%) was quantified from (n=3/group, 6 fields/sample). (D) Immunoblots of pAKT in liver lysatesfrom shGFP and shAdipsin-treated mice. Right: Densitometric analysis of relative pAKT protein levels. (E–H) Quantification of body weight, liver weight, epididymal adipose tissue depot, and total plasma triglyceride levels in shGFP and shAdipsin-treated mice. (I) mRNA expression of gluconeogenesis genes (Creb1, Fbp1, Fbp2, G6pc1, G6pc3, Mdh1, Mdh2, Pck1, Pck2, Pfkb1, Ppara) from shGFP and shAdipsin treated mice. (J) Immunoblots of PCK1 in liver lysatesfrom shGFP and shAdipsin-treated mice. Right: Densitometric analysis of relative PCK1protein levels (n=3/group). Data are presented as mean ± SD. ** P < 0.01; statistical significance was determined using an unpaired two-tailed t-test. ns, not significant.

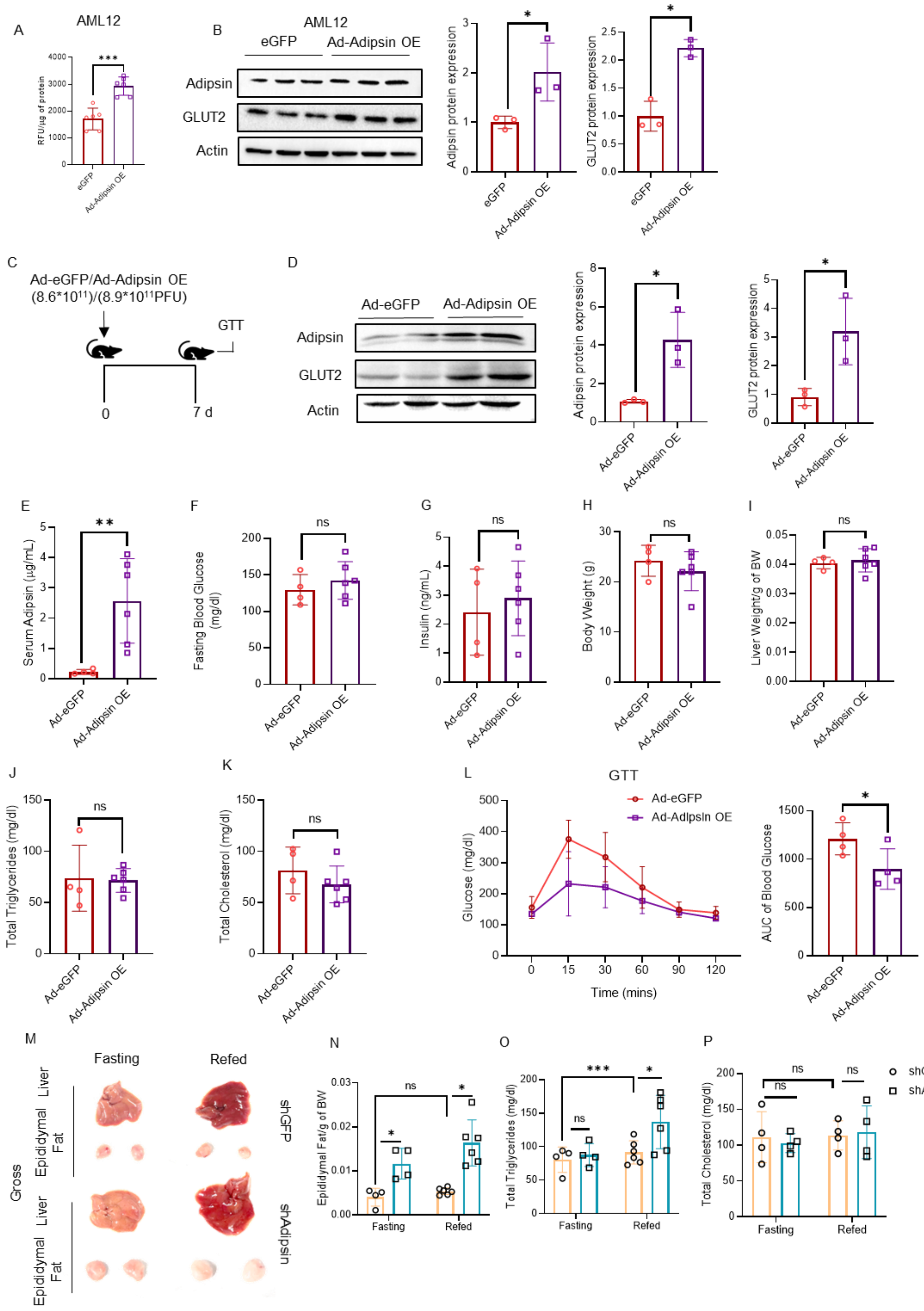


**Supplementary Figure 4. Hepatic adipsin overexpression improves glycemic control.** (A) Glucose uptake assay in AML12 cells following adenovirus-mediated overexpression (Ad-Adipsin OE) of Adipsin. (B) Immunoblot analysis of adipsin and GLUT2 protein homogenates following adenovirus-mediated overexpression (Ad-Adipsin OE) of Adipsin in AML12 cells. Right panel: Densitometric analysis of Adipsin and GLUT2 (n=3/group). (C) Schematic representation of adenovirus-mediated hepatic adipsin overexpression (Ad-Adipsin OE). (D) Immunoblot analysis of Adipsin and GLUT2 protein levels in liver lysates from Ad-eGFP- and Ad-Adipsin OE-treated mice (n=3/group). (E–K) Serum adipsin levels (E), fasting blood glucose (F), insulin levels (G), body weight (H), liver weight (I), total triglycerides (J), and cholesterol levels (K) in Ad-eGFP and Ad-Adipsin OE treated mice. (L) Glucose Tolerance Test (GTT) in Ad-eGFP- and Ad-Adipsin OE-treated mice. (M) Representative gross liver and epididymal adipose tissue depotin fasting and refed mice. (N) Epididymal adipose tissue weight. (O, P) Total cholesterol (O), and triglyceride levels (P) in fasting and refed groups of shGFP and shAdipsin-treated mice. Mice n=8 (Ad-eGFP), n=8 (Ad-Adipsin OE). Data are presented as mean ± SD. ** $P < 0.01$; statistical significance was determined using an unpaired two-tailed t-test. ns, not significant.
